# The draft chromosome-level genome assembly of tetraploid ground cherry (*Prunus fruticosa* Pall.) from long reads

**DOI:** 10.1101/2021.06.01.446499

**Authors:** Thomas W. Wöhner, Ofere F. Emeriewen, Alexander H.J. Wittenberg, Harrie Schneiders, Ilse Vrijenhoek, Júlia Halász, Károly Hrotkó, Katharina J. Hoff, Lars Gabriel, Jens Keilwagen, Thomas Berner, Mirko Schuster, Andreas Peil, Jens Wünsche, Stephan Kropop, Henryk Flachowsky

**Affiliations:** Julius Kühn Institute (JKI) – Federal Research Centre for Cultivated Plants, Institute for Breeding Research on Fruit Crops, Pillnitzer Platz 3a, D-01326, Dresden, Germany; Keygene N.V., P.O. Box 216, 6700 AE Wageningen, Netherlands; Department of Genetics and Plant Breeding, Faculty of Horticultural Science, Szent István University, Ménesi Str. 44, Budapest 1118, Hungary; Department of Floriculture and Dendrology, Institute of Landscape Architecture, Urban Planning and Ornamental Horticulture, Hungarian University of Agriculture and Life Science, Villányi Str. 35-43, Budapest 1118, Hungary; Institute of Mathematics and Computer Science, University of Greifswald, Walther-Rathenau-Str. 47, 17489 Greifswald, Germany; Julius Kühn Institute (JKI) – Federal Research Centre for Cultivated Plants, Institute for Biosafety in Plant Biotechnology, Erwin-Baur-Str. 27, D-06484 Quedlinburg, Germany; University of Hohenheim, Institute of Special Crops and Crop Physiology, 70593 Stuttgart, Germany; Center for Functional Genomics of Microbes, University of Greifswald, Felix-Hausdorff-Str. 8, 17489 Greifswald, Germany; Jagdweg 1-3, 01159 Dresden, Germany

**Keywords:** genome assembly, long read, *P. fruticosa*, ground cherry, tetraploid

## Abstract

**Background:** Cherries are stone fruits and belong to the economically important plant family of *Rosaceae* with worldwide cultivation of different species. The ground cherry, *Prunus fruticosa* Pall. is one ancestor of cultivated sour cherry, an important tetraploid cherry species. Here, we present a long read chromosome-level draft genome assembly and related plastid sequences using the Oxford Nanopore Technology PromethION platform and R10.3 pore type.

**Finding:** The final assemblies obtained from 117.3 Gb cleaned reads representing 97x coverage of expected 1.2 Gb tetraploid (2n=4x=32) and 0.3 Gb haploid (1n=8) genome sequence of *P. fruticosa* were calculated. The N50 contig length ranged between 0.3 and 0.5 Mb with the longest contig being ∼6 Mb. BUSCO estimated a completeness between 98.7 % for the 4n and 96.1 % for the 1n datasets.

Using a homology and reference based scaffolding method, we generated a final consensus genome sequence of 366 Mb comprising eight chromosomes. The N50 scaffold was ∼44 Mb with the longest chromosome being 66.5 Mb.

The repeat content was estimated to ∼190 Mb (52 %) and 58,880 protein-coding genes were annotated. The chloroplast and mitochondrial genomes were 158,217 bp and 383,281 bp long, which is in accordance with previously published plastid sequences.

**Conclusion:** This is the first report of the genome of ground cherry (*P. fruticosa*) sequenced by long read technology only. The datasets obtained from this study provide a foundation for future breeding, molecular and evolutionary analysis in *Prunus* studies.

## Data Description

### Context

Cherries are stone fruits belonging to the important family of *Rosaceae* fruit crops, which are produced for fresh fruit consumption or industrial processing [1]. The worldwide production of cherries was 4 million metric tons on an area of 6.7 million ha [2] in 2019. Nevertheless, cherry production worldwide is threaten by changing climatic conditions, which promote pests, e.g., *Drosophila suzukii* and *Rhagoletis cerasi*, diseases, e.g., *Monilinia laxa* and *Blumeriella jaapii*, as well as unfavourable abiotic conditions, e.g., hail or late frost [1, 3]. Breeding of new cultivars that are resistant to biotic stress factors and adapted to local climate conditions could contribute to sustainable cultivation in the long-term and secure future production. Donors for breeding and introgression of new characters and traits can be found in wild/related species of the genus *Prunus* [4–6]. The ground cherry (*Prunus fruticosa* Pall.) is a wild *Prunus* species with a small shrub-like habitus that is native from middle Europe to Western Siberia and Western China [7, 8]. The natural habitats vary from open landscapes with steppe characteristics, the edges of open forests [9–11] or hillsides with stony soils [12]. *Prunus fruticosa* is a self-incompatible [13] tetraploid (2n=4x=32) species with an estimated genome size of 1.31 pg determined by flow cytometry analysis [14]. It is the progenitor of sour cherry (*P. cerasus* L.), which developed by natural hybridization from unreduced pollen of sweet cherry (*P. avium* L.) with *P.* fruticosa [15, 16]. *Prunus fruticosa* is a valuable genetic resource for breeding of varieties adapted to drought and low temperatures [17, 18] because of its growth at cold and semi-arid sites and its edible fruits [7]. Due to its dwarf habitus, the species has been used as a donor for cherry rootstock breeding in several programmes [19–21]. Like other *Rosaceae* fruit species, cherries are perennial crops and breeding of new cultivars is labour intensive and time consuming [22]. Genome sequencing advances breeding processes enormously by providing insights into evolution and comparative studies with related species, determining the positions of putative genes, which may control different traits, and allowing for the possibility for marker-assisted selection. Hence several genomes of other *Prunus* species [23–30] as well as other members of the *Rosaceae* family [31–33] have been sequenced in recent years. The sizes of *Prunus* genomes so far sequenced range between 250-300 Mbp with high synteny of the eight basic chromosomes [3]. However, sequencing and assembling plant genomes is still a challenging task. Although the commercialization of third-generation sequencing technology has enabled rapid generation of giga-bases of data, most genome sequences are still fragmented or incomplete due to size, composition and structure (repeat content) of genomes with many reference genomes presented as drafts. The availability of long read sequencing technologies can solve these problems and offers many more advantages [34].

In this study, we present a draft assembly of the *P. fruticosa* Pall. genome generated with long read Oxford Nanopore Technology (ONT). Using the final assembly for reference based scaffolding, eight chromosome scale pseudomolecules were constructed and subsequently used for gene annotation. This data provides additional information, which may be useful for breeding and genetic diversity studies in cherry and the genus *Prunus* in general.

## Material and Methods

### Plant Material, DNA extraction and ONT sequencing

*Prunus fruticosa* Pall. young leaf material (tetraploid, short type, size ca. 30-50 cm) was collected in its natural habitat [8] from a single tree (in situ) in Budapest, Hármashatárhegy (Fig. 1, coordinates 47°33’15.322’’N, 18°59’49.623’’E). Snap frozen plant material was sent to the sequencing service provider KeyGene N.V. (Wageningen, The Netherlands) for high molecular weight DNA extraction, purification and nanopore sequencing analysis. High molecular weight DNA was extracted by KeyGene N.V. using nuclei isolated from frozen leaves ground under liquid nitrogen, as described elsewhere [35, 36]. Genomic DNA was quality controlled with a Qubit device (Thermo Fisher Scientific, Waltham, MA, USA) and length was determined using the Femto Pulse instrument (Agilent, California). Short DNA fragments were removed using the Circulomics SRE XL kit (Circulomics, Baltimore, MD, USA) following the manufactures instruction. Finally 2 µg AMPure purified genomic DNA per flow cell (AMPure PB, Pacific Biosciences, California) was used as input for library construction using the 1D Genomic DNA ligation SQK_LSK110 library prep kit (Oxford Nanopore Technologies, Oxford, UK). Subsequently, the library was loaded on three PromethION FLO PRO003 (R10.3 pore, early access pore) flow cells and run on PromethION P24 platform according to the manufacturer’s recommendations. Basecalling was performed in real-time on the compute module (PromethION version: 20.06.9/Guppy4.0.11). Only passed reads with a Q-value threshold of seven were used for further data analysis.

**Figure 1.**
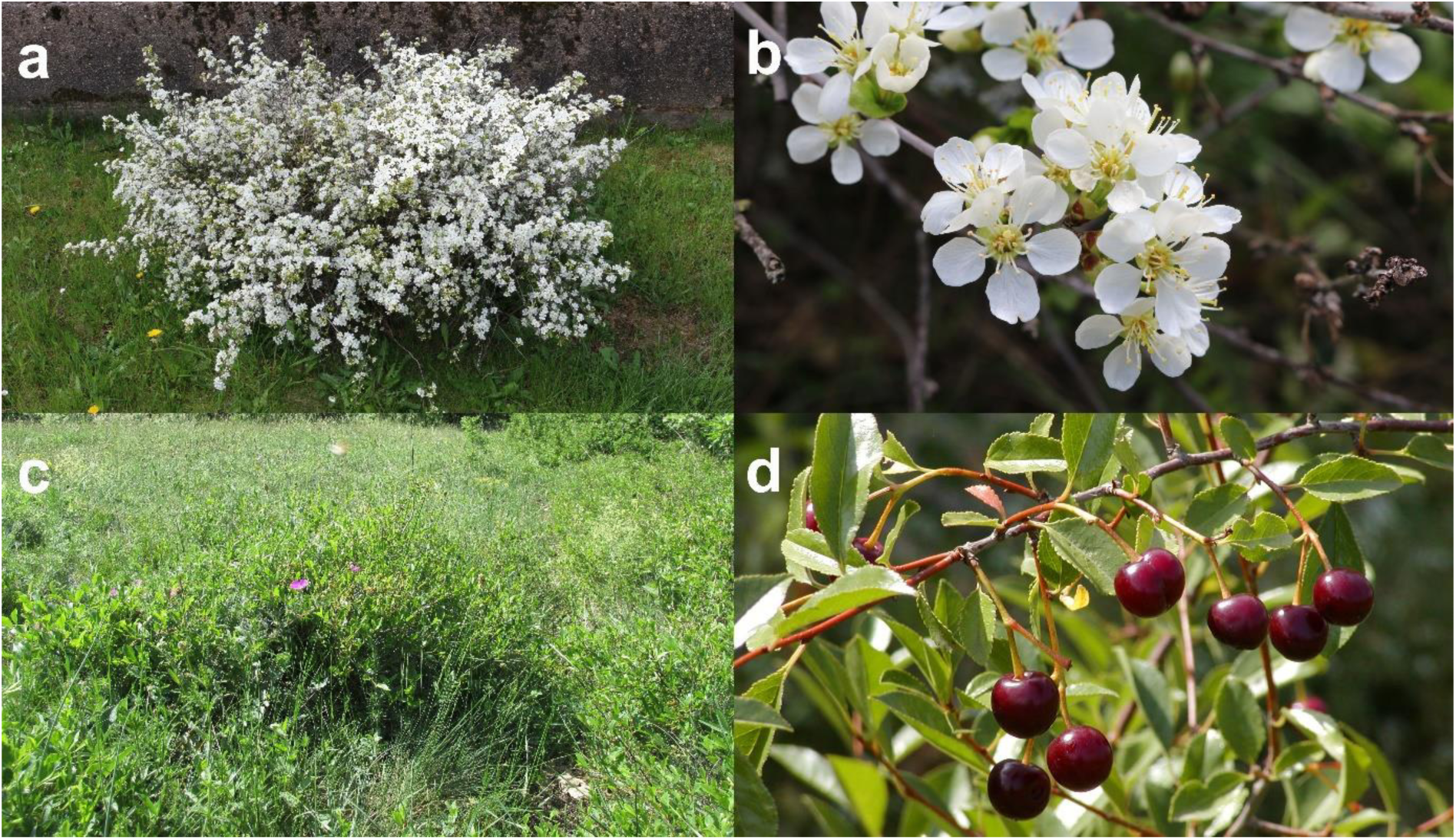
Morphology of *P. fruticosa* Pall.. (a) flowering habitus, (b) inflorescence, (c) mature shrub in the natural habitat in Hungary and (d) leafs and fruits.

### De novo assembly and scaffolding

Raw data assembly was performed using a combination of the aligner Minimap2 (2.16-r922) and the assembler Miniasm (0.2-r137-dirty) using a 20x, 30x and 50x coverage/length cut-offs at default settings. Three runs of Racon (v1.4.10) subsequently improved base accuracy of the interim contig assembly using a 10 Kb length cut-off and one run of Medaka (1.01) using all raw reads for consensus calling. The sequences of the obtained contig assembly were collapsed with two runs of Purge Dups (V1.0.1) using default settings. The BUSCO (Benchmark Universal Single-Copy Orthologs - Galaxy Version 4.1.4) software was used for quantitative and quality assessment of the genome assemblies based on near-universal single-copy orthologs. The genome sequence of *P. avium* ‘Tieton’ ([37], GenBank assembly accession: GCA_014155035.1) was used as a matrix for reference guided scaffolding of the final assembly (purged2) using RAGOO (v1.11) with the standard settings [38]. Final sequence statistics were calculated with CLC Mainworkbench (v20.0.4). The generated *P. fruticosa* genome (Pf_1.0) was hard masked with NCBI WindowMasker [39] implementation on the CoGe platform [40]. Synteny comparisons between *P. avium* ‘Tieton’ and *P. persica* ‘Lovell’ ([24], GenBank assembly accession: GCA_000346465.2) with Pf_1.0 were performed with SynMap2 [41] using the standard program settings.

### Annotation

A species-specific repeat library for Pf_1.0 was first generated with RepeatModeler 1.0.11 [42]. The obtained dataset was then used for repetitive sequence identifcation and masking in Pf_1.0 with ReapeatMasker 4.0.7 [43]. As no RNA-seq data for *P. fruticosa* was available, publicly available RNA-seq data [44] from the close relative *P. cerasus* ‘Schattenmorelle’ (SRR2290965) was downloaded from NCBI and mapped to Pf_1.0 using HISAT2 2.1.0 [45].

The structural gene annotation of genomic features is result of a combination of ab initio and homology-based gene annotation. Ab initio gene prediction was performed with both BRAKER1 [46, 47] and BRAKER2 [48]. The BRAKER pipeline in general leverages extrinsic data, such as spliced alignments from short read RNA-Seq or large-scale protein to genome alignments for executing self-training GeneMark-ET/EP [49] [50, 51]with help of SAMtools [52], and BamTools [53], or GeneMark-EP+ [54], with DIAMOND [55], GeneMark-ES [56], and Spaln2 [57, 58] for generating an evidence-supported training gene set for the gene finder AUGUSTUS. AUGUSTUS then predicts genes with evidence where available [59] and in *ab initio* mode in local absence of evidence [60]. OrthoDB v.10 *Plantae* partition [61] and related species proteins [*P. armeniaca* (GCA_903112645.1)*, P. persica* (GCF_000346465.2), *P. mume* (GCF_000346735.1), *P. dulcis* (GCF_902201215.1) and *P. avium* (GCF_002207925.1)] obtained from GenBank were used as reference protein dataset for BRAKER2. Gene predictions from BRAKER1 and BRAKER2 were combined into one transcript set by filtering the union of transcripts from both predictions in context with their support by the evidence generated with PrEvCo v. 0.1.0 (https://github.com/LarsGab/PrEvCo). The obtained *ab initio* annotation was augmented with additional GFF attributes using the GeMoMa module AnnotationEvidence.

Homology-based gene annotation was performed with GeMoMa version 1.7.2beta [62] using the mapped RNA-seq data from ‘Schattenmorelle’ and the genome and gene annotation from the following reference organisms that are available at NCBI: *A. thaliana* (TAIR10.1, RefSeq GCF_000001735.4)*, M. domestica* (GDDH1, GCF_002114115.1)*, F. vesca* (FraVesHawai_1.0, GCF_000184155.1)*, P. avium* (PAV_r1.0, GCF_002207925.1)*, P. persica* (Prunus_persica_NCBIv2, GCF_000346465.2)*, P. mume* (P.mume_V1.0, GCF_000346735.1)*, P. dulcis* (ALMONDv2, GCF_902201215.1) and *P. armeniaca* (pruArmRojPasHapCUR, GCA_903112645.1).

The augmented *ab initio* gene annotation from BRAKER and the eight homology-based gene predictions from GeMoMa were combined using the GeMoMa module GAF yielding a final gene annotation. BUSCO with set embryophyta_odb10 (Galaxy Version 4.1.4) was used for the assessment of protein completeness. For handling alternative transcripts correctly and not as duplicates, a custom script was ran on the BUSCO full table, assigning gene ID instead of transcript ID. The functional annotation was performed with the obtained protein files using InterproScan at Galaxy Europe using default parameters [63–65] and [66].

Noncoding RNA prediction was performed with tRNAscan (Galaxy version 0.4), Aragorn (Galaxy version 0.6), barrnap (Galaxy version 1.2.1) and INFERNAL (cmsearch with rFAM 11.0, Galaxy Version 1.1.2.0).

The chloroplast and mitochondria sequences were annotated with GeSeq [67] using the references for chloroplast from *P. fruticosa* (GenBank accession MT916286) published by [68] and mitochondria from *P. avium* (GenBank accession MK816392) published by [69]. GeSeq pipeline analysis was performed using the annotation packages ARAGORN, blatN, blatX, Chloe and HMMER.

## Data validation and quality control

We report the use of Oxford Nanopore technology to assemble a high-quality reference genome of *P*. *fruticosa* – the first report in a tetraploid *Prunus* species. Previously described genomes in *Prunus* applied Illumina, PacBio or shotgun sequencing techniques [25, 26, 29]. However, Wang et al. [28] reported a combination of Oxford Nanopore and Illumina technologies for sweet cherry. Table S1 summarizes the assembly statistics of our study. We generated 4.5 million raw reads (124.7 Gb), which is considerably lower compared to the read output of *P*. *avium* cultivars [25, 28]. After cleaning, approximately 4.0 million reads comprised 117.3 Gb in total (mean q = 9.96), which were generated by the R10.3 PromethION flow cells representing ∼97x coverage of the estimated tetraploid genome size of 1.2 Gb. Compared to Wang et al. [28], the R10.3 flow cells produced longer reads with higher quality (Table S1). A mean of 1,347,740 (SD = 135.304) reads with a N50 length of 41,236 (SD = 275) bp and 39.1 (SD = 4.2) Gb per flow cell were obtained (Table S1). Based on the results of the raw data assemblies (Table S2), it was decided to continue with the obtained 30x coverage Miniasm assembly with a length cut-off at 62.3 kb. After three runs of Racon and one run of Medaka consensus calling, the final assembly covered approximately four times the estimated haploid genome size of ∼0.3 Gb, indicating we were able to separate the parental haplotypes (4n) to a large extent. Consensus calling resulted in a total assembly size of 1161.5 Mb, represented by 4.426 contigs with an N50 contig size of 325 Kb and the longest contig almost 5,9 Mb (Table 1). Two runs of Purge Dups were performed to collapse the haplotype-separated assembly in order to reduce the duplicated content to a haplotype consensus sequence (1n). The purged_2x assembly data set has a size of 376,7 Mb and consists 1.275 contigs with an N50 contig size of 533.426 bp. This assembly was used as input for reference-guided scaffolding using RaGoo and the genome sequence of *P. avium* ‘Tieton’ [28]. The obtained sequence file consists of nine scaffolds representing eight chromosomes and one sequence with concatenated unmapped data (unassigned). The final *Prunus fruticosa* 1.0 genome sequence (Fig. 2) consists of 366,5 Mb with a N50 size of scaffolds about 43,818.497 bp and G+C of 37.74 %, A+T of 62.22 % and only 0.03 % gaps (N). The longest scaffold is 66,497,422 bp (Table 2). Compared to the genome sequences available so far in *Prunus* [24, 25, 28], the genome of *P. fruticosa* is the most complete obtained from long read sequencing only.

**Figure 2.**
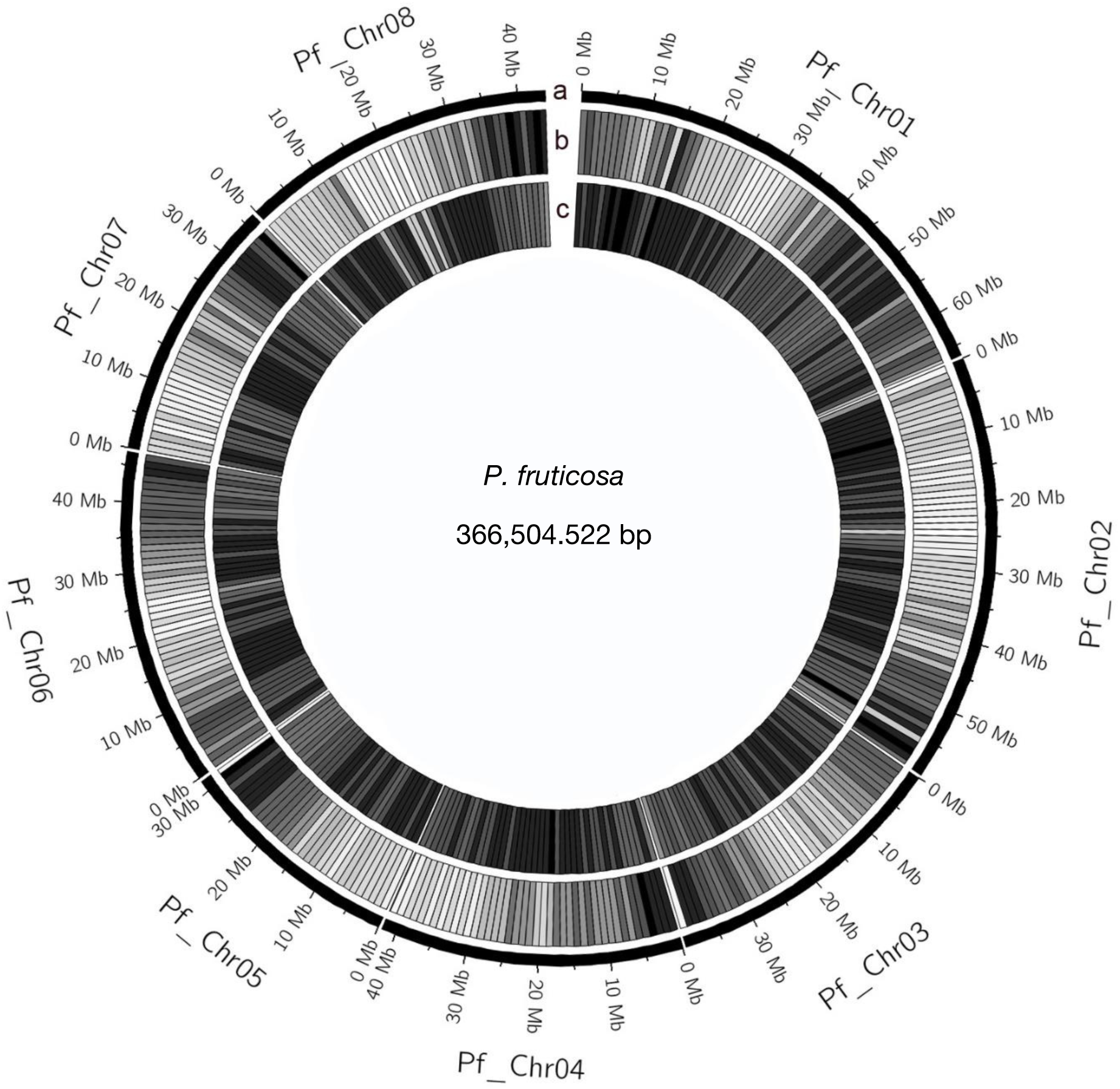
The genome of *P. fruticosa.* Circos plot of the 8 pseudomolecules. (a) Chromosome length (Mb); (b) gene density in blocks of 1 MB; (c) repeat density in blocks of 1 Mb.

**Table 1.** Statistics of different datasets a<u>nd assemblie</u>s from *P. fruticosa*

| Data set / assembly | Ploidy | Number of contigs | Contig N50 (bp) | Longest contig (bp) | Total contig length (Mb) |
| --- | --- | --- | --- | --- | --- |
| All reads | 4n | 4,525.811 | 40.963 | 1,257.508 | 1247,375 |
| Passed reads | 4n | 4,043.192 | 41.244 | 732.658 | 1172,679 |
| Miniasm/Minimap | 4n | 4.399 | 324.889 | 5,840.253 | 1147,459 |
| Racon | 4n | 4.381 | 326.739 | 5,954.545 | 1161,2 |
| Medaka | 4n | 4.426 | 325.453 | 5,956.772 | 1161,5 |
| Purge dups_1x | 1n | 1.516 | 501.505 | 5,956.772 | 480,6 |
| Purge dups_2x | 1n | 1.275 | 533.462 | 5,956.772 | 376,7 |

**Table 2.**
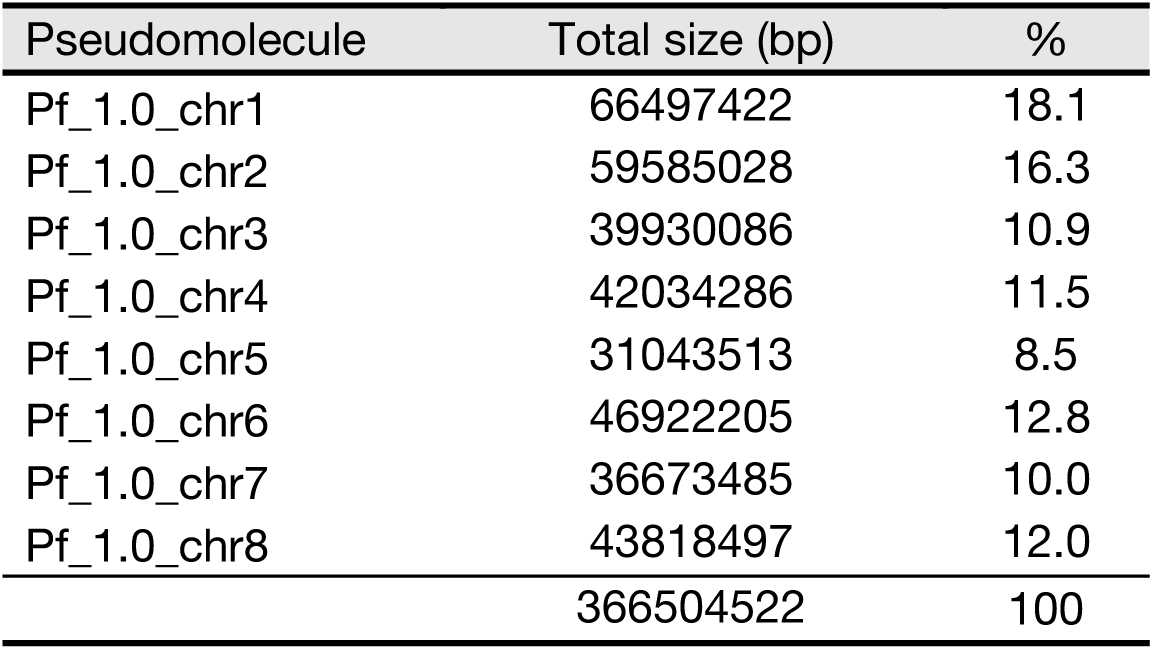
Pseudomolecule statistics for Pf_1.0

BUSCO analysis resulted in 98.6 % - 98.7 % completeness for the representing 4n Racon and Medaka generated data sets. The comparison of BUSCO results (Fig. 3) on assembly completeness between the Racon only and the Racon and Medaka data sets (Table 1) indicates that consensus generation by Medaka increases the number of duplicated genes (from 89.7 % to 92.4 %) and improves the consensus accuracy. The obtained assembly sequences (1n) after haplotig removal showed a decrease of duplicated BUSCOS (from 92.4 % to 12.5 %) and an increase of single BUSCOS (from 6.3 % to 83.6 %). *P. fruticosa* 1.0 results outlined in Figure 3 show a 96.4 % completeness. Compared to the genome sequence of *P. persica* (99.3 %) and *P. avium* (98.3 %) which represent the highest genome completeness of published datasets, the obtained long read only assemblies (98.7 %) and consensus genome sequence (96.4 %) from this study shows a comparably high genome completeness.

**Figure 3.**
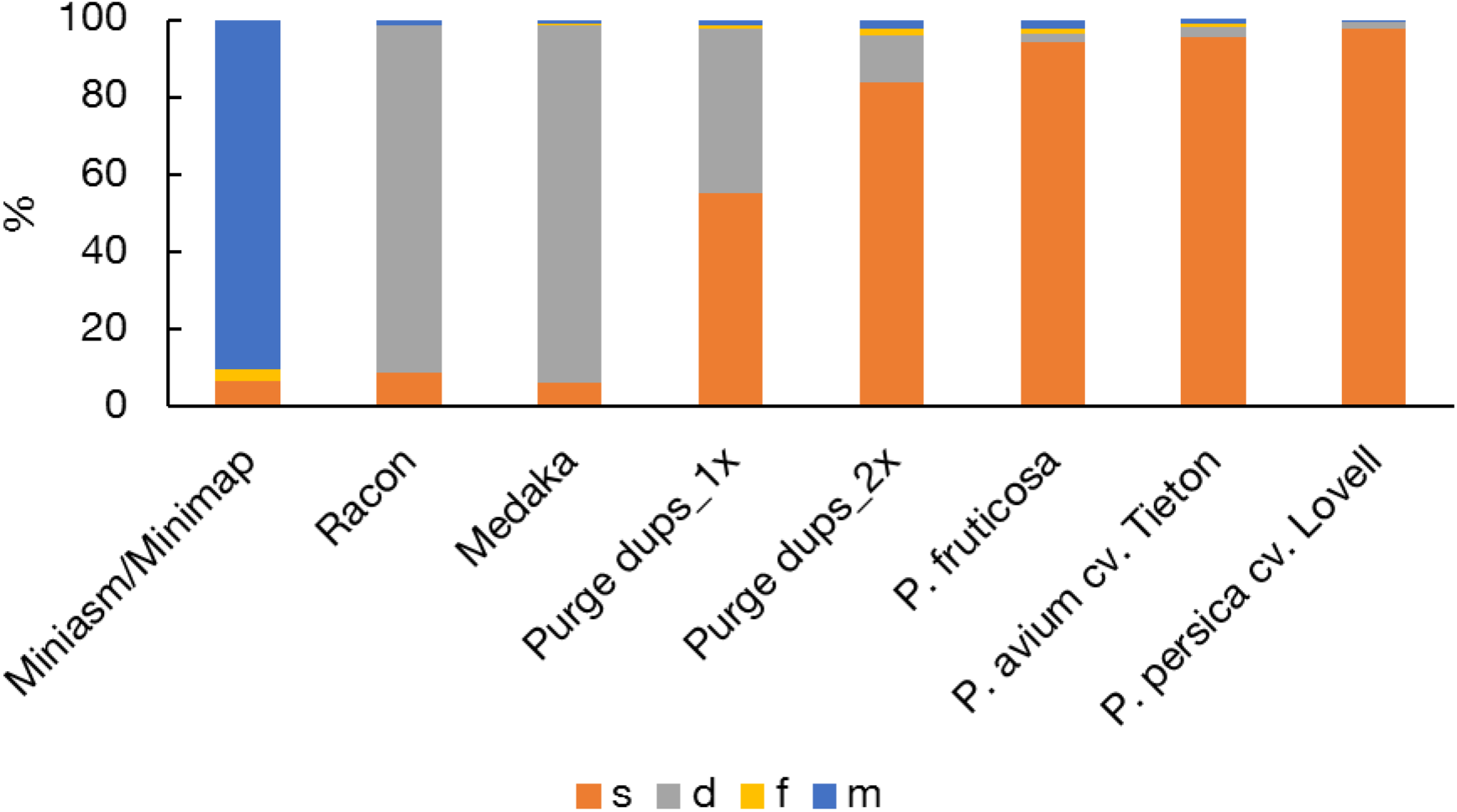
Analysis of completeness of different *P. fruticosa* datasets compared to *P. avium* cv. Tieton and *P. persica* cv. Lovell by mapping of a set of universal single-copy orthologs using BUSCO. The bar charts indicate complete **<u>s</u>**ingle copy (orange), complete **<u>d</u>**uplicated (gray), **f**ragmented (yellow) and **<u>m</u>**issing (blue) genes. For evaluation the embryophyta_odb10 BUSCO dataset (n=1614) was used. *P. fruticosa* 1.0 show a 96.4 % completeness (S: 94.1 %, D: 2.3 %, F: 1.3 %, M: 2.3 %, n: 1614) which almost reaches the completeness of *P. avium* cv. ‘Tieton’ (C: 98.3 %, S: 95.6 %, D: 2.7 %, F: 0.5 %, M:1.5 %, n:1614) and *P. persica* ‘Lovell’ (C: 99.3 %, S: 97.5 %, D: 1.8%, F: 0.1 %, M: 0.6 %, n:1614).

Our approach detected 189,7 Mb of repetitive sequences (51.75 % of the genome) and 42,1 Mb (11.5 %) unknown elements. Repetitive sequences observed in other *Prunus* species [25–27, 29, 33] ranged from 37.1% in *P. persica* [50] to 59.4 % in *P. avium* [28]. However, similar to *P*. *avium* [25], the repeated sequences observed in our study comprised mainly of the class (I) LTR Gypsy retrotransposons and *Copia*. LTR was the most abundant element in our findings with 20.88 % followed by Copia with 7.59 % (Table 3).

**Table 3.**
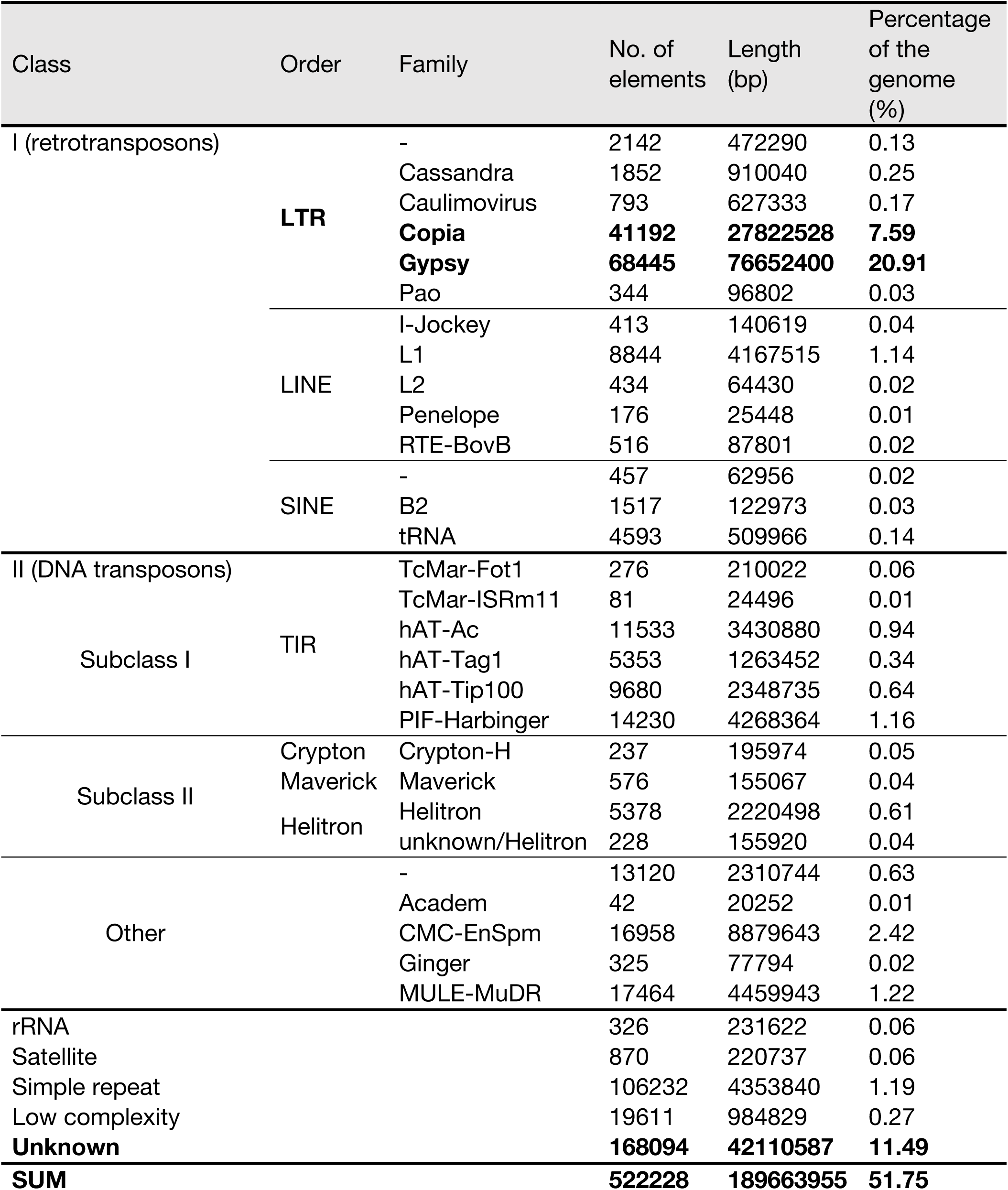
Characterization of repetitive sequences of *P. fruticosa* 1.0

We employed similar strategy as reported elsewhere namely homology-based, *de novo* and transcriptome supported approaches [28, 37] to call repeats, predict protein-coding genes and perform functional annotation. Using RNA-Seq data from *P. cerasus* ‘Schattenmorelle’ [44] and the augmented gene predictions from BRAKER with eight homology-based gene predictions from GeMoMa we predicted 58.880 protein-coding transcripts representing 84.524 orthologs within Pf_1.0 with a mean length of 3.580 bp and a mean protein length of 355 aa (Table 4). The number of protein-coding transcripts was considerably larger in this study than 38.275 predicted for *P. avium* ‘Tieton’ [28] and 43.349 transcripts predicted in *P. avium* ‘Satonishiki’ [25]. A total of 86.7 % (75,113) proteins was functionally annotated by InterproScan resulting in 852.470 annotated protein domains and sites from 15 protein databases (Table 4). A total of 2.301 (Aragorn) and 2.559 (tRNA scan) tRNA and 576 rRNA sequences were detected. Infernal search reveals 36.757 consensus RNA secondary structure profiles. BUSCO analysis for transcriptome completeness (embryophyta_odb10 dataset) reveals 1,552 (96.2 %) complete (81.8 % single and complete, 14.4 % duplicated and complete) and 62 (3.8 %) fragmented (1.7 %) or missing (2.1 %) BUSCOs (Fig. 4).

**Figure 4.**
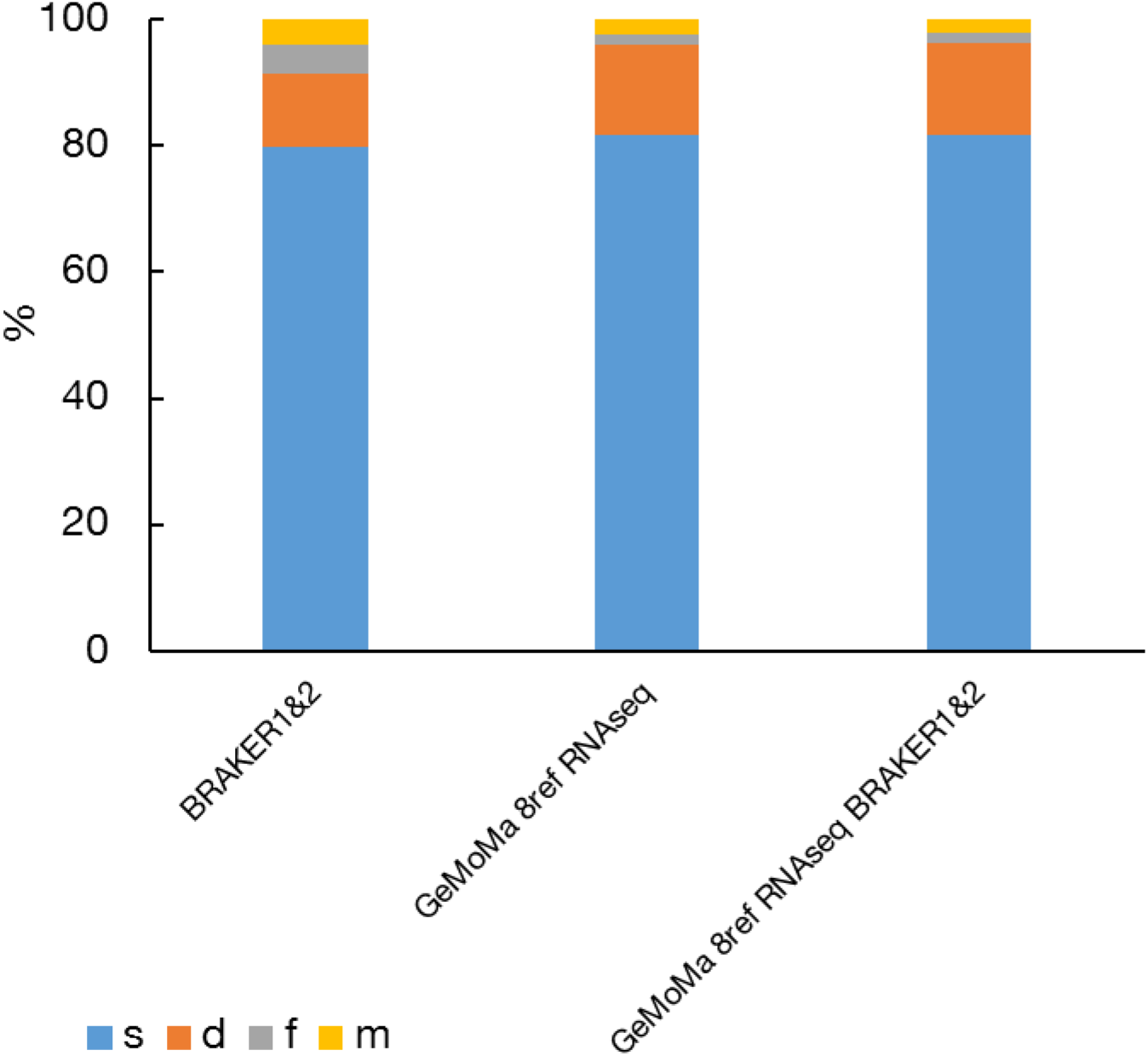
Analysis of completeness of different protein sets obtained with different structural annotation strategies. The bar charts indicate complete **<u>s</u>**ingle copy (orange), complete **<u>d</u>**uplicated (gray), **f**ragmented (yellow) and **<u>m</u>**issing (blue) genes. For evaluation the embryophyta_odb10 BUSCO dataset (n=1614) was used.

**Table 4.** Functional annotation results generated by interproscan using BRAKER & GeMoMa combination of ab-initio and homology-based structural gene annotation and statistics

| Interproscan annotations | No. |
| --- | --- |
| Coils | 14627 |
| Gene3D | 82428 |
| Hamap | 1336 |
| PANTHER | 150554 |
| Pfam | 95569 |
| Phobius | 197895 |
| PIRSF | 5075 |
| PRINTS | 48332 |
| ProSitePatterns | 19050 |
| ProSiteProfiles | 52557 |
| SignalP_EUK | 7914 |
| SMART | 42825 |
| SUPERFAMILY | 64033 |
| TIGRFAM | 10603 |
| TMHMM | 59672 |
| Sum | **852470** |

| Transcripts | No. | % |
| --- | --- | --- |
| total | 58880 |  |
| orthologs | 84524 | 100 |
| annotated | 73315 | 86.7 |
| annotated GO | 45196 | 53.5 |
| annotated pathways | 5247 | 6.2 |
| domains | 62431 | 73.9 |
| Mean length (bp) | 3580 |  |
| Mean length of predicted proteins | 355 |  |

The obtained chloroplast genome sequence (Fig. 5a) was 158,130 bp long (GC 36.6 %) with a typical quadripartite structure consisting a large (86,242 bp) and a small (19,143) single-copy region and two inverted repeats (IRA 26,372 bp, IRB 26,373 bp). The GC contents of each region were 34.1 % (LSC), 30.1 % (SSC) and 42.5 % for IRA and IRB each. The size, the structure and the GC content values are similar to those reported previously for the chloroplast genome of *P. fruticosa* (Yang et al. 2020). Forty-five tRNA (ARAGORN), eight rRNA (each with HMMER and blatN) and 116 protein-coding genes (HMMER) were annotated.

**Figure 5.**
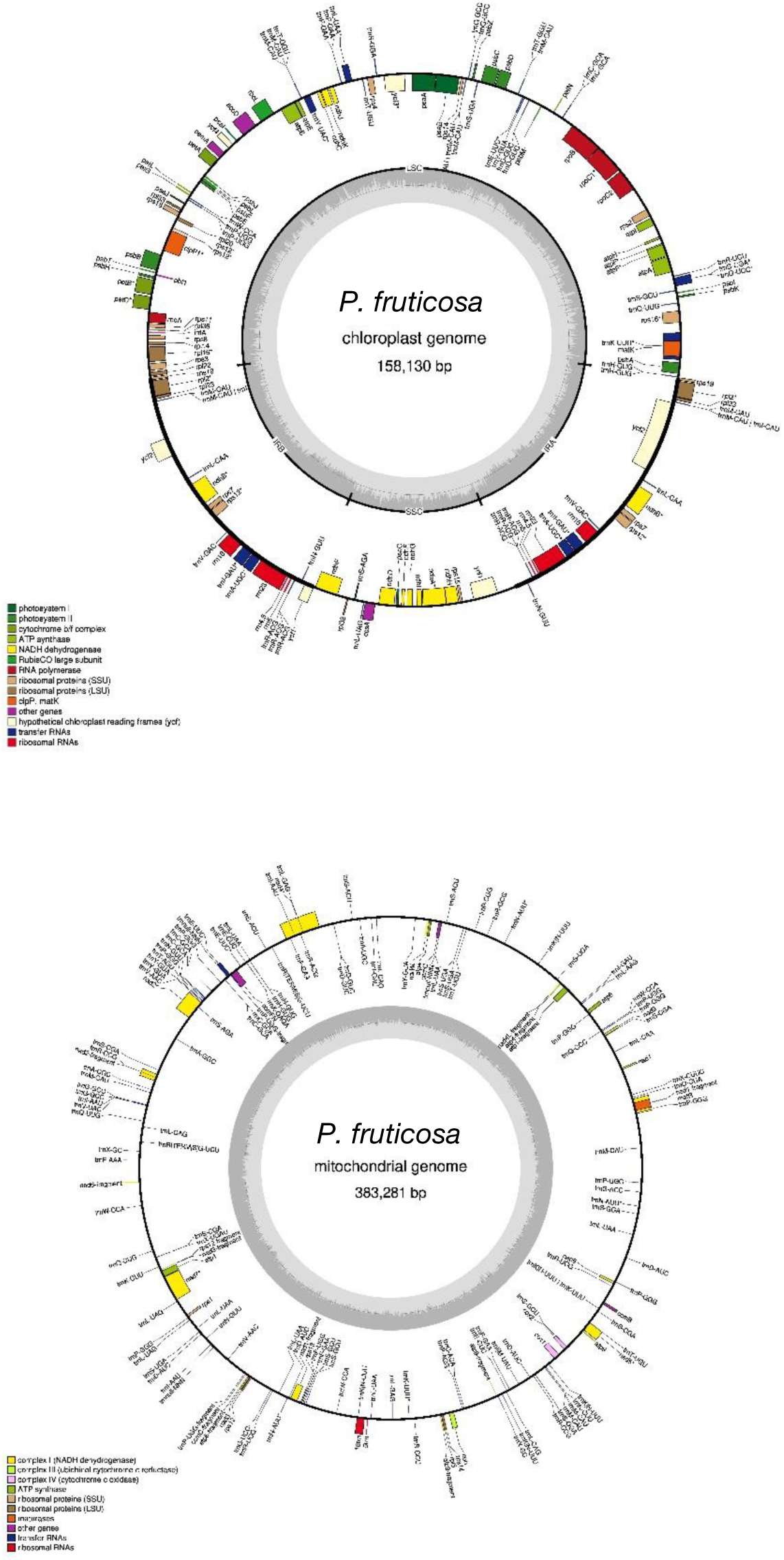
The chloroplast (a) and mitochondrial (b) genome sequence of *P. fruticosa* 1.0 obtained from the contigs utg000088l and utg001396I in the medaka assembly sequence. Annotation was performed using GeSeq (Tillich et al. 2017).

We present for the first time a mitochondrial genome for *P. fruticosa* (Fig. 5b) with a length of 383,281 bp and a GC content of 45.7 %. The results of the mitochondria genome is similar to the mitochondria genome of *P. avium* ‘Summit’ [69] where a total of 68 protein coding genes, including 27 tRNA (ARAGORN) and two rRNA (blatN) were annotated.

We compared sequence synteny between *P. fruticosa* and *P. persica* and *P. fruticosa* and *P. avium* (Fig. 6). The synteny analysis involved at least two transcripts of annotated genes in each representative genome (Fig. 6a). As indicated in Table S3, a higher percentage of transcripts (77.5 % to 87.3 %) were mapped between the homologues chromosomes from *P. persica* and Pf_1.0 compared to the transcripts from *P. avium* (72.1% to 56.3 %). In general, the assembled genome of *P. fruticosa* shows a good synteny with the genomes of *P. persica* [24] and *P.* avium [28]. Figure 6b shows the synteny analysis using masked sequences (i.e. without repetitive sequences). The results obtained confirm strong synteny between the compared genomes and strongly suggest the high quality of the obtained genome sequence.

**Figure 6.**
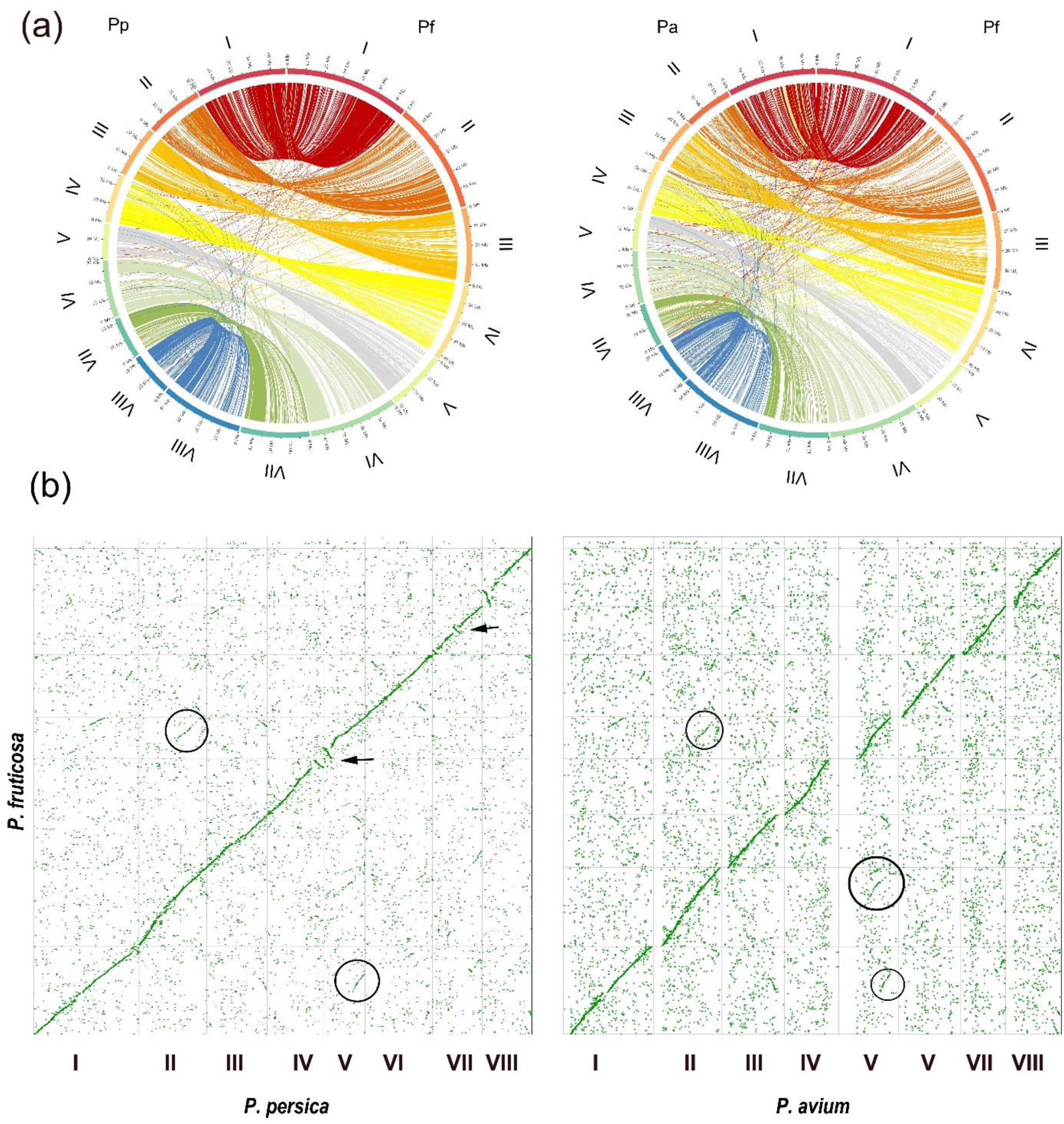
Synteny between *P. fruticosa*, *P. persica* ‘Lovell’ and *P. avium* ‘Tieton’. (a) Circos plots showing transcripts of *P. persica* (Pp, left) and *P. avium* (Pa, right) annotated in *P. fruticosa* (Pf). Each string represents at least two transcripts in a 50k bp cluster. (b) Syntenic dot plot of the nucleotide sequences between *P. fruticosa, P. persica* and *P. avium.* Before plotting, the sequences were hard masked by the NCBI window maker implication on the CoGe webpage. Several inversions (arrows) and out-paralogs (circles) were identified between the sequences.

## Re-use potential

For the first time, we report a draft genome scale-assembly of tetraploid *Prunus* species. This was achieved using Nanopore sequencing technology, confirming that this technology alone can sufficiently produce a high-quality genome without additional sequencing using Illumina [70]. This genome will be valuable in exploiting genetic information for breeding programs; will enhance our understanding of genetics of this species relative to breeding as well as molecular and evolutionary analysis in the genus *Prunus*.

## Data Availability

Data supporting the findings of this study are deposited into the Open Agrar repository [71] and on personal request to the corresponding author. The ground cherry genome has been submitted to NCBI and is available after review.

## Authors’ contribution

TW, OE wrote the manuscript. AW, HS and IV performed DNA isolation, sequencing and genome assembly. JH and KH provided the plant material. KH, JK, LG and TB performed annotation of the dataset. SK performed the scaffolding and TW the did the interproscan and synteny analysis. HF, JW, MS and AP conceived the study and made substantial contributions to its design, acquisition, analysis and interpretation of data. All authors contributed equally to the finalization of the manuscript.

## Supplemental

**Table S1.** Statistics of three different datasets for *P. fruticosa* generated with R10.3 PromethION cells (passed reads)

| Data set | Ploidy | Number of reads | Total gigabases (Gb) | N50 length (bp) | Mean length (bp) | Max length (bp) | Mean q |
| --- | --- | --- | --- | --- | --- | --- | --- |
| 200917_PAF21731 | 4n | 1,498.775 | 43.7 | 41.257 | 29.104 | 732.658 | 9.9 |
| 200922_PAF21408 | 4n | 1,306.840 | 38.2 | 41.499 | 29.251 | 529.628 | 10 |
| 200922_PAF21416 | 4n | 1,237.604 | 35.4 | 40.951 | 28.615 | 624.468 | 10 |
|  | Sum | 4,043.219 | 117.3 |  |  |  |  |
|  | Mean | 1,347.740 | 39.1 | 41.236 | 28.990 | 628.918 | 9.96 |
|  | SD | 135.304 | 4.2 | 275 | 333 | 101.588 | 0.06 |

**Table S2.** Assembly statistics of tetraploid *P. fruticosa*

| x-coverage* | 20x |  | 30x |  | 50x |  |
| --- | --- | --- | --- | --- | --- | --- |
| Assembly | MM** + 3x racon | + medaka | MM** + 3x racon | + medaka | MM** + 3x racon | + medaka |
| Final no. of seq. | 3.413 | 3.460 | **4.381** | **4.426** | <u>4.662</u> | <u>4.727</u> |
| Seq. length<br>characters (bp) |  |  |  |  |  |  |
| Cutoff | 69.042 |  | 63.011 |  | 55.002 |  |
| Total | 989.839.499 | 988.573.324 | **1.162.237.634** | **1.161.456.281** | 1.212.434.570 | 1.211.504.091 |
| Mean | 290.020 | 285.715 | **265.290** | **262.417** | 260.067 | 256.294 |
| SD | 244.496 | 242.372 | 263.004 | 262.317 | 277.742 | 276.312 |
| Min | 4.077 | 149 | 2.605 | 73 | 4.077 | 99 |
| Max | 3.012.684 | 3.006.345 | 5.954.545 | 5.956.772 | 6.443.975 | 6.446.220 |
| N25 | 661.833 | 654.351 | 598.365 | 598.625 | 596.874 | 596.453 |
| N50 | 368.933 | 367.276 | **326.739** | **325.453** | 314.773 | 313.883 |
| N75 | 204.724 | 204.105 | 186.200 | 185.890 | 182.574 | 181.440 |
| N90 | 144.538 | 144.375 | 135.082 | 134.607 | 133.209 | 132.581 |

\* fold coverage based on the estimation that the haploid genome size is ~300 Mbp
\*\* MM – Miniasm & Minimap

**Table S3a.**
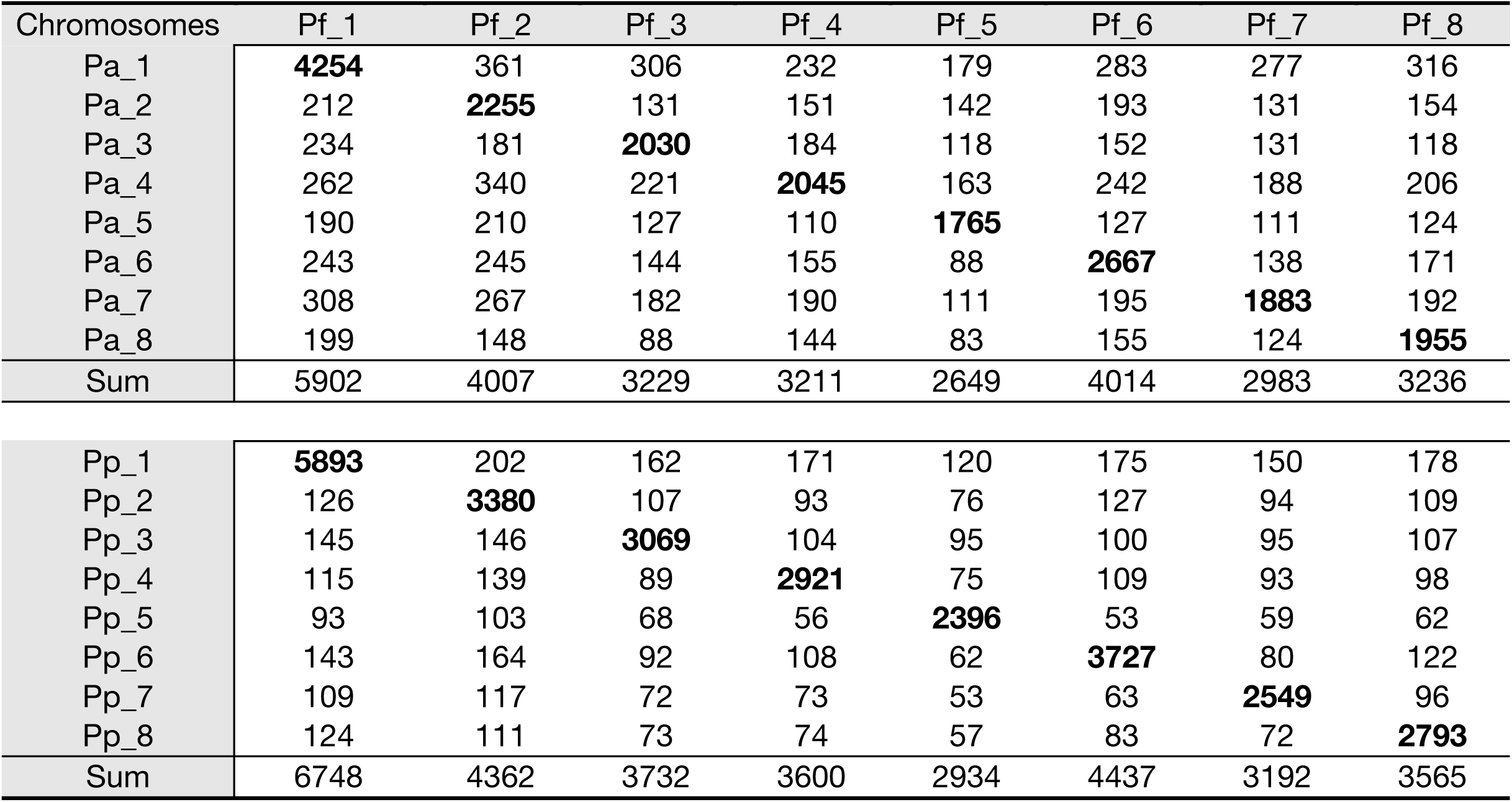
Matrix of shared number of transcripts annotated from *P. avium* (Pa) and *P. persica* (Pp) to *P. fruticosa* (Pf) Pf_1.0 within each chromosomes

| Chromosomes | Pf_1 | Pf_2 | Pf_3 | Pf_4 | Pf_5 | Pf_6 | Pf_7 | Pf_8 |
| --- | --- | --- | --- | --- | --- | --- | --- | --- |
| Pa_1 | **4254** | 361 | 306 | 232 | 179 | 283 | 277 | 316 |
| Pa_2 | 212 | **2255** | 131 | 151 | 142 | 193 | 131 | 154 |
| Pa_3 | 234 | 181 | **2030** | 184 | 118 | 152 | 131 | 118 |
| Pa_4 | 262 | 340 | 221 | **2045** | 163 | 242 | 188 | 206 |
| Pa_5 | 190 | 210 | 127 | 110 | **1765** | 127 | 111 | 124 |
| Pa_6 | 243 | 245 | 144 | 155 | 88 | **2667** | 138 | 171 |
| Pa_7 | 308 | 267 | 182 | 190 | 111 | 195 | **1883** | 192 |
| Pa_8 | 199 | 148 | 88 | 144 | 83 | 155 | 124 | **1955** |
| Sum | 5902 | 4007 | 3229 | 3211 | 2649 | 4014 | 2983 | 3236 |
| Pp_1 | **5893** | 202 | 162 | 171 | 120 | 175 | 150 | 178 |
| Pp_2 | 126 | **3380** | 107 | 93 | 76 | 127 | 94 | 109 |
| Pp_3 | 145 | 146 | **3069** | 104 | 95 | 100 | 95 | 107 |
| Pp_4 | 115 | 139 | 89 | **2921** | 75 | 109 | 93 | 98 |
| Pp_5 | 93 | 103 | 68 | 56 | **2396** | 53 | 59 | 62 |
| Pp_6 | 143 | 164 | 92 | 108 | 62 | **3727** | 80 | 122 |
| Pp_7 | 109 | 117 | 72 | 73 | 53 | 63 | **2549** | 96 |
| Pp_8 | 124 | 111 | 73 | 74 | 57 | 83 | 72 | **2793** |
| Sum | 6748 | 4362 | 3732 | 3600 | 2934 | 4437 | 3192 | 3565 |

**Table S3b.** Matrix of shared number of transcript in percent annotated from *P. avium* (Pa) and *P. persica* (Pp) to *P. fruticosa* (Pf) Pf_1.0 within each chromosomes

| Chromosomes | Pf_1 | Pf_2 | Pf_3 | Pf_4 | Pf_5 | Pf_6 | Pf_7 | Pf_8 |
| --- | --- | --- | --- | --- | --- | --- | --- | --- |
| Pa_1 | **72.08** | 9.01 | 9.48 | 7.23 | 6.76 | 7.05 | 9.29 | 9.77 |
| Pa_2 | 3.59 | **56.28** | 4.06 | 4.70 | 5.36 | 4.81 | 4.39 | 4.76 |
| Pa_3 | 3.96 | 4.52 | **62.87** | 5.73 | 4.45 | 3.79 | 4.39 | 3.65 |
| Pa_4 | 4.44 | 8.49 | 6.84 | **63.69** | 6.15 | 6.03 | 6.30 | 6.37 |
| Pa_5 | 3.22 | 5.24 | 3.93 | 3.43 | **66.63** | 3.16 | 3.72 | 3.83 |
| Pa_6 | 4.12 | 6.11 | 4.46 | 4.83 | 3.32 | **66.44** | 4.63 | 5.28 |
| Pa_7 | 5.22 | 6.66 | 5.64 | 5.92 | 4.19 | 4.86 | **63.12** | 5.93 |
| Pa_8 | 3.37 | 3.69 | 2.73 | 4.48 | 3.13 | 3.86 | 4.16 | **60.41** |
| Pp_1 | **87.33** | 4.63 | 4.34 | 4.75 | 4.09 | 3.94 | 4.70 | 4.99 |
| Pp_2 | 1.87 | **77.49** | 2.87 | 2.58 | 2.59 | 2.86 | 2.94 | 3.06 |
| Pp_3 | 2.15 | 3.35 | **82.23** | 2.89 | 3.24 | 2.25 | 2.98 | 3.00 |
| Pp_4 | 1.70 | 3.19 | 2.38 | **81.14** | 2.56 | 2.46 | 2.91 | 2.75 |
| Pp_5 | 1.38 | 2.36 | 1.82 | 1.56 | **81.66** | 1.19 | 1.85 | 1.74 |
| Pp_6 | 2.12 | 3.76 | 2.47 | 3.00 | 2.11 | **84.00** | 2.51 | 3.42 |
| Pp_7 | 1.62 | 2.68 | 1.93 | 2.03 | 1.81 | 1.42 | **79.86** | 2.69 |
| Pp_8 | 1.84 | 2.54 | 1.96 | 2.06 | 1.94 | 1.87 | 2.26 | **78.35** |

## References

1. Quero-Garcia J, Iezzoni A, Lopez-Ortega G, Peace C, Fouche M, Dirlewanger E, Schuster M. Advances and challenges in cherry breeding 2019: Burleigh Dodds Science Publishing.

2. FAO. Crops. 2021. http://www.fao.org/faostat/en/#data/QC/. Accessed 17 Feb 2021.

3. Aranzana MJ, Decroocq V, Dirlewanger E, Eduardo I, Gao ZS, Gasic K, et al. Prunus genetics and applications after de novo genome sequencing: achievements and prospects. Horticulture research. 2019;6:1–25.

4. Schuster M, Grafe C, Hoberg E, Schütze W. Interspecific hybridization in sweet and sour cherry breeding. Acta horticulturae. 2013.

5. Wen J, Berggren ST, Lee C-H, Ickert-Bond S, Yi T-S, Yoo K-O, et al. Phylogenetic inferences in Prunus (Rosaceae) using chloroplast ndhF and nuclear ribosomal ITS sequences. J Syst Evol. 2008;46:322–32.

6. Stegmeir T, Schuster M, Sebolt A, Rosyara U, Sundin GW, Iezzoni A. Cherry leaf spot resistance in cherry (Prunus) is associated with a quantitative trait locus on linkage group 4 inherited from P. canescens. Molecular breeding. 2014;34:927–35.

7. Faust M, Surányi D. Origin and dissemination of cherry. Horticultural Reviews. 1997;19:263–317.

8. Hrotkó K, Feng Y, Halász J. Spontaneous hybrids of Prunus fruticosa Pall. in Hungary. Genetic Resources and Crop Evolution. 2020;67:489–502.

9. Jäger EJ, Seidel D. Unterfamilie Prunoideae. Hegi Illustrierte Flora von Mitteleuropa. 1995;4.

10. Meusel H, Jäger EJ, Weinert E. Vergleichende chorologie der zentraleuropaischen flora. 1965.

11. Macková L, Vít P, Urfus T. Crop-to-wild hybridization in cherries—Empirical evidence from Prunus fruticosa. Evolutionary applications. 2018;11:1748–59.

12. Mratinić E, Kojić M. Wild fruit species of Serbia. Agricultural Research Institute of Serbia, Belgrade. 1998.

13. Pruski K. Tissue culture propagation of Mongolian cherry (Prunus fruticosa L.) and Nanking cherry (Prunus tomentosa L.). In: Protocols for micropropagation of woody trees and fruits: Springer; 2007. p. 391–407.

14. Macková L, Vít P, Ďurišová Ľ, Eliáš P, Urfus T. Hybridization success is largely limited to homoploid Prunus hybrids: a multidisciplinary approach. Plant Systematics and Evolution. 2017;303:481–95.

15. Olden EJ, Nybom N. On the origin of Prunus cerasus L. Hereditas. 1968;59:327–45.

16. Brettin TS, Karle R, Crowe EL, af Iezzoni. Chloroplast Inheritance and DNA Variation in Sweet, Sour, and Ground Cherry. The Journal of heredity. 2000;91:75–9.

17. Mitschurin IW. Ausgewählte Schriften 1951. In: Verlag Kultur und Fortschritt, editor. Ausgewählte Schriften 1951.

18. Bors RH. Dwarf sour cherry breeding at the University of Saskatchewan. Acta horticulturae. 2005.

19. Cummins JN. Vegetatively propagated selections of Prunus fruticosa as dwarfing stocks for cherry. Fruit Var Hort Dig. 1972.

20. Plock H. Bedeutung der Prunus fruticosa Pall. als Zwergunterlage fur Suss-und Sauerkirschen. Mitt Rebe Wein Obstbau Fruchteverwert. 1973.

21. Hein K. Zwischenbericht über eine Prüfung der Steppenkirsche (P. fruticosa) und anderen Süsskirchenunterlagen und Unterlagenkombinationen. Erwerbsobstbau. 1979;21:219.

22. Peace C, Norelli J. Genomics approaches to crop improvement in the Rosaceae. In: Genetics and genomics of Rosaceae: Springer; 2009. p. 19–53.

23. Ahmad R, Parfitt DE, Fass J, Ogundiwin E, Dhingra A, Gradziel TM, et al. Whole genome sequencing of peach (Prunus persica L.) for SNP identification and selection. BMC genomics. 2011;12:1–7.

24. Verde I, Jenkins J, Dondini L, Micali S, Pagliarani G, Vendramin E, et al. The Peach v2. 0 release: high-resolution linkage mapping and deep resequencing improve chromosome-scale assembly and contiguity. BMC genomics. 2017;18:1–18.

25. Shirasawa K, Isuzugawa K, Ikenaga M, Saito Y, Yamamoto T, Hirakawa H, Isobe S. The genome sequence of sweet cherry (Prunus avium) for use in genomics-assisted breeding. DNA Research. 2017;24:499–508.

26. Baek S, Choi K, Kim G-B, Yu H-J, Cho A, Jang H, et al. Draft genome sequence of wild Prunus yedoensis reveals massive inter-specific hybridization between sympatric flowering cherries. Genome biology. 2018;19:1–17.

27. Shirasawa K, Esumi T, Hirakawa H, Tanaka H, Itai A, Ghelfi A, et al. Phased genome sequence of an interspecific hybrid flowering cherry,‘Somei-Yoshino’(Cerasus× yedoensis). DNA Research. 2019;26:379–89.

28. Wang J, Liu W, Zhu D, Hong P, Zhang S, Xiao S, et al. Chromosome-scale genome assembly of sweet cherry (Prunus avium L.) cv. Tieton obtained using long-read and Hi-C sequencing. Horticulture research. 2020;7:1–11.

29. Zhang Q, Chen W, Sun L, Zhao F, Huang B, Yang W, et al. The genome of Prunus mume. Nature communications. 2012;3:1–8.

30. Callahan AM, Zhebentyayeva TN, Humann JL, Saski CA, Galimba KD, Georgi LL, et al. Defining the‘HoneySweet’insertion event utilizing NextGen sequencing and a de novo genome assembly of plum (Prunus domestica). Horticulture research;8:8.

31. Velasco R, Zharkikh A, Affourtit J, Dhingra A, Cestaro A, Kalyanaraman A, et al. The genome of the domesticated apple (Malus× domestica Borkh.). Nature genetics. 2010;42:833–9.

32. Daccord N, Celton J-M, Linsmith G, Becker C, Choisne N, Schijlen E, et al. High-quality de novo assembly of the apple genome and methylome dynamics of early fruit development. Nature genetics. 2017;49:1099–106.

33. Jiang F, Zhang J, Wang S, Yang L, Luo Y, Gao S, et al. The apricot (Prunus armeniaca L.) genome elucidates Rosaceae evolution and beta-carotenoid synthesis. Horticulture research. 2019;6:1–12.

34. Jain M, Olsen HE, Paten B, Akeson M. The Oxford Nanopore MinION: delivery of nanopore sequencing to the genomics community. Genome biology. 2016;17:1–11.

35. Zhang M, Zhang Y, Scheuring CF, Wu C-C, Dong JJ, Zhang H-B. Preparation of megabase-sized DNA from a variety of organisms using the nuclei method for advanced genomics research. nature protocols. 2012;7:467–78.

36. Datema E, Hulzink RJM, Blommers L, Valle-Inclan JE, van Orsouw N, Wittenberg AHj, Vos M de. The megabase-sized fungal genome of Rhizoctonia solani assembled from nanopore reads only. BioRxiv. 2016:84772.

37. Liu C, Feng C, Peng W, Hao J, Wang J, Pan J, He Y. Chromosome-level draft genome of a diploid plum (Prunus salicina). GigaScience. 2020;9:giaa130.

38. Alonge M, Soyk S, Ramakrishnan S, Wang X, Goodwin S, Sedlazeck FJ, et al. RaGOO: fast and accurate reference-guided scaffolding of draft genomes. Genome biology. 2019;20:1–17.

39. Morgulis A, Gertz EM, Schäffer AA, Agarwala R. WindowMasker: window-based masker for sequenced genomes. Bioinformatics. 2006;22:134–41.

40. Lyons EH. CoGe, a new kind of comparative genomics platform: Insights into the evolution of plant genomes: University of California, Berkeley; 2008.

41. Haug-Baltzell A, Stephens SA, Davey S, Scheidegger CE, Lyons E. SynMap2 and SynMap3D: web-based whole-genome synteny browsers. Bioinformatics. 2017;33:2197–8.

42. Smit AF, Hubley R, Green P. RepeatModeler Open-1.0. 2008–2015. Seattle, USA: Institute for Systems Biology. Available from: httpwww. repeatmasker. org, Last Accessed May. 2015;1:2018.

43. Smit AF, Hubley R, Green P. RepeatMasker Open-4.0. 2013–2015 2015.

44. Jo Y, Chu H, Cho JK, Choi H, Lian S, Cho WK. De novo transcriptome assembly of a sour cherry cultivar, Schattenmorelle. Genomics data. 2015;6:271–2.

45. Kim D, Paggi JM, Park C, Bennett C, Salzberg SL. Graph-based genome alignment and genotyping with HISAT2 and HISAT-genotype. Nature biotechnology. 2019;37:907–15.

46. Hoff KJ, Lange S, Lomsadze A, Borodovsky M, Stanke M. BRAKER1: unsupervised RNA-Seq-based genome annotation with GeneMark-ET and AUGUSTUS. Bioinformatics. 2016;32:767–9.

47. Hoff KJ, Lomsadze A, Borodovsky M, Stanke M. Whole-genome annotation with BRAKER. In: Gene prediction: Springer; 2019. p. 65–95.

48. Brůna T, Hoff KJ, Lomsadze A, Stanke M, Borodovsky M. BRAKER2: Automatic eukaryotic genome annotation with GeneMark-EP+ and AUGUSTUS supported by a protein database. NAR Genomics and Bioinformatics. 2021;3:lqaa108.

49. Lomsadze A, Burns PD, Borodovsky M. Integration of mapped RNA-Seq reads into automatic training of eukaryotic gene finding algorithm. Nucleic acids research. 2014;42:e119–e119.

50. Verde I, Abbott AG, Scalabrin S, Jung S, Shu S, Marroni F, et al. The high-quality draft genome of peach (Prunus persica) identifies unique patterns of genetic diversity, domestication and genome evolution. Nature genetics. 2013;45:487–94.

51. Verde I, Abbott AG, Scalabrin S, Jung S, Shu S, Marroni F, et al. The high-quality draft genome of peach (Prunus persica) identifies unique patterns of genetic diversity, domestication and genome evolution. Nature genetics. 2013;45:487–94.

52. Li H, Handsaker B, Wysoker A, Fennell T, Ruan J, Homer N, et al. The sequence alignment/map format and SAMtools. Bioinformatics. 2009;25:2078–9.

53. Barnett DW, Garrison EK, Quinlan AR, Strömberg MP, Marth GT. BamTools: a C++ API and toolkit for analyzing and managing BAM files. Bioinformatics. 2011;27:1691–2.

54. Brůna T, Lomsadze A, Borodovsky M. GeneMark-EP+: Eukaryotic Gene Prediction With Self-Training in the Space of Genes and Proteins. NAR Genomics and Bioinformatics. 2020;2:lqaa026.

55. Buchfink B, Xie C, Huson DH. Fast and Sensitive Protein Alignment Using DIAMOND. Nature methods. 2015;12:59–60.

56. Lomsadze A, Ter-Hovhannisyan V, Chernoff YO, Borodovsky M. Gene identification in novel eukaryotic genomes by self-training algorithm. Nucleic acids research. 2005;33:6494–506.

57. Iwata H, Gotoh O. Benchmarking spliced alignment programs including Spaln2, an extended version of Spaln that incorporates additional species-specific features. Nucleic acids research. 2012;40:e161–e161.

58. Gotoh O, Morita M, Nelson DR. Assessment and refinement of eukaryotic gene structure prediction with gene-structure-aware multiple protein sequence alignment. Bmc Bioinformatics. 2014;15:1–13.

59. Stanke M, Diekhans M, Baertsch R, Haussler D. Using native and syntenically mapped cDNA alignments to improve de novo gene finding. Bioinformatics. 2008;24:637–44.

60. Stanke M, Schöffmann O, Morgenstern B, Waack S. Gene prediction in eukaryotes with a generalized hidden Markov model that uses hints from external sources. Bmc Bioinformatics. 2006;7:1–11.

61. Kriventseva EV, Kuznetsov D, Tegenfeldt F, Manni M, Dias R, Simão FA, Zdobnov EM. OrthoDB v10: sampling the diversity of animal, plant, fungal, protist, bacterial and viral genomes for evolutionary and functional annotations of orthologs. Nucleic acids research. 2019;47:D807–D811.

62. Keilwagen J, Hartung F, Grau J. GeMoMa: Homology-based gene prediction utilizing intron position conservation and RNA-seq data. In: Gene Prediction: Springer; 2019. p. 161–177.

63. Cock PJA, Grüning BA, Paszkiewicz K, Pritchard L. Galaxy tools and workflows for sequence analysis with applications in molecular plant pathology. PeerJ. 2013;1:e167.

64. Zdobnov EM, Apweiler R. InterProScan: protein domains identifier. Bioinformatics (Oxford, England). 2001;17:847–8.

65. Quevillon E, Silventoinen V, Pillai S, Harte N, Mulder N, Apweiler R, Lopez R. InterProScan: protein domains identifier. Nucleic acids research. 2005;33:W116–W120.

66. Hunter S, Apweiler R, Attwood TK, Bairoch A, Bateman A, Binns D, et al. InterPro: the integrative protein signature database. Nucleic acids research. 2009;37:D211–D215.

67. Tillich M, Lehwark P, Pellizzer T, Ulbricht-Jones ES, Fischer A, Bock R, Greiner S. GeSeq–versatile and accurate annotation of organelle genomes. Nucleic acids research. 2017;45:W6–W11.

68. Yang Y-X, Tian M-H, Liu X-H, Li Y, Sun Z-S. Complete chloroplast genome of Prunus fruticosa and its implications for the phylogenetic position within Prunus sensulato (Rosaceae). Mitochondrial DNA Part B. 2020;5:3624–6.

69. Yan M, Zhang X, Zhao X, Yuan Z. The complete mitochondrial genome sequence of sweet cherry (Prunus avium cv.‘summit’). Mitochondrial DNA Part B. 2019;4:1996–7.

70. Huang Y-T, Liu P-Y, Shih P-W. High-Quality Genomes of Nanopore Sequencing by Homologous Polishing. BioRxiv. 2020.

71. Wöhner TW, Emeriewen OF, Wittenberg AHj, Schneiders H, Vrijenhoek I, Halász J, et al. Supporting materials for - The draft chromosome-level genome assembly of tetraploid ground cherry (Prunus fruticosa Pall.) from long reads. 2021. https://www.openagrar.de/receive/openagrar_mods_00070329. Accessed 1 Jun 2021.

